# The effect of sonic hedgehog on motor neuron positioning in the spinal cord during chicken embryo development

**DOI:** 10.1101/178491

**Authors:** Ciqing Yang, Xiaoying Li, Qiuling Li, Qiong Li, Han Li, Bichao Zhang, Juntang Lin

## Abstract

Sonic hedgehog (Shh) is a vertebrate homologue of the secreted *Drosophila* protein hedgehog, and is expressed by the notochord and the floor plate in the developing spinal cord. Shh provides signals relevant for positional information, cell proliferation, and possibly cell survival depending on the time and location of the expression. Although the role of Shh in providing positional information in the neural tube has been experimentally proven, the exact underlying mechanism still remains unclear. In this study, we report that overexpression of Shh affects motor neuron positioning in the spinal cord during chicken embryo development by inducing abnormalities in the structure of the motor column and motor neuron integration. In addition, Shh overexpression inhibits the expression of dorsal transcription factors and commissural axon projections. Our results indicate that correct location of Shh expression is the key to the formation of the motor column. In conclusion, the overexpression of Shh in the spinal cord not only affects the positioning of motor neurons, but also induces abnormalities in the structure of the motor column.

## INTRODUCTION

During central nervous system development, many factors can be controlled to ensure normal development. The early embryonic vertebrate neural tube consists of proliferating progenitors and terminally differentiating neurons with a defined distribution pattern (Cayuso et al., 2006). The notochord and floor plate at the ventral midline of the neural tube determine, in part, the organization of the developing spinal cord (Pringle et al., 1996). These structures also emit signals that can induce the development of distant motor neurons (Yamada et al., 1991, 1993; Tanabe et al., 1995). In the ventral spinal cord, motor neurons (MN) are grouped in motor columns according to their identity and their target muscle (Luxey et al., 2015). Different motor neurons express different sets of transcription factors. For instance, HB9 is expressed in all somatic MN, whereas Foxp1, Lim1, and Islet1 are all expressed in lateral motor column MN at high levels (Vermot et al., 2005; Bonanomi and Pfaff, 2010; Santiago et al., 2014; Luxey et al., 2015). All these transcription factors have been shown to contribute to the establishment of MN organization in the spinal cord. Indeed, gain and loss of function of HB9, Islet1, Islet2, Lim1, and Foxp1 lead to important defects of MN positioning within the spinal cord during embryo development (Kania et al., 2000; Odden et al., 2002; Bréjot et al., 2006; Hutchinson and Eisen, 2006; Rousso et al., 2008; Otaegi et al., 2011). Although the role of these transcription factors in MN positioning in the spinal cord is well established, little is known regarding their potential effector genes (Luxey et al., 2015).

Sonic hedgehog (Shh) is a vertebrate homologue of the secreted protein encoded by the *Drosophila* gene hedgehog (Lee et al., 1992; Nusslein-Volhard et al., 1980), and is expressed by the notochord and floor plate at the time when these structures exert their inductive activities (Riddle et al., 1993; Echelard et al., 1993). In the central nervous system, Shh plays an important role in ventral specification along the entire neural axis. In ventral regions, this protein acts as a long-range graded signal that controls the pattern of neurogenesis (Jessell, 2000; Briscoe and Ericson, 2001). Misexpression of Shh in vertebrate embryos can induce the differentiation of floor plate cells at ectopic locations in the neural tube (Echelard et al., 1993; Krauss et al., 1993; Roelink et al., 1994). Shh provides signals relevant to positional information, cell proliferation, and possibly cell survival depending on the timing and location of the expression (Riddle et al., 1993; Peterson et al., 2012; Yang et al., 2015). Although the role of Shh in providing positional information in the neural tube has been experimentally established, the mechanism underlying this phenomenon remains unclear.

In this study, we focus on the role of Shh in motor neuron positioning in the spinal cord during chicken embryo development by inducing its misexpression in the embryonic spinal cord. We examined the gene expression in Shh-transfected spinal cord and followed spinal cord development. Shh expression can directly or indirectly affect the development of multiple structures. Moreover, the localization of dorsal-ventral cell types was determined to analyze the effects of Shh in cell type specification. The results of these studies indicated that Shh affects the expression of dorsal transcription factors Pax3 and Pax7 and the positioning of ventral motor neurons in the spinal cord.

## RESULTS

### Shh overexpression in the developing chicken spinal cord

In ovo electroporation, a technique by which the plasmid can be unilaterally electroporated, was performed to examine the role of Shh in the developing spinal cord. Two experimental groups were designed as follows: 1) electroporation of pCAGGS-GFP (0.25 μg/μL) – control group, 2) co-electroporation of pCAGGS-Shh (4 μg/μL) + pCAGGS-GFP (0.25 μg/μL) - experimental group. Electroporation was performed on the chicken embryonic spinal cord at stage 17 (E2.5). After 36, 60, and 84 h following electroporation, GFP-positive embryos were collected at stage 24-26 (E4-E6), and the overexpression of Shh was clearly observed using in situ hybridization (Fig. 1A-C, arrows [→] indicate the areas of Shh overexpression). To control for individual differences, data from the same spinal cord, where the transfected and non-transfected sides served as experimental and control tissue, respectively, were matched. Shh was expressed by the notochord and floor plate in the developing chicken spinal cord (Fig. 1D-F). As the notochord is also known to induce differentiation of other ventral cell types within the neural tube, including motor neurons, it can be hypothesized that Shh produced by the notochord may be required for motor neuron differentiation.

**Fig. 1.**
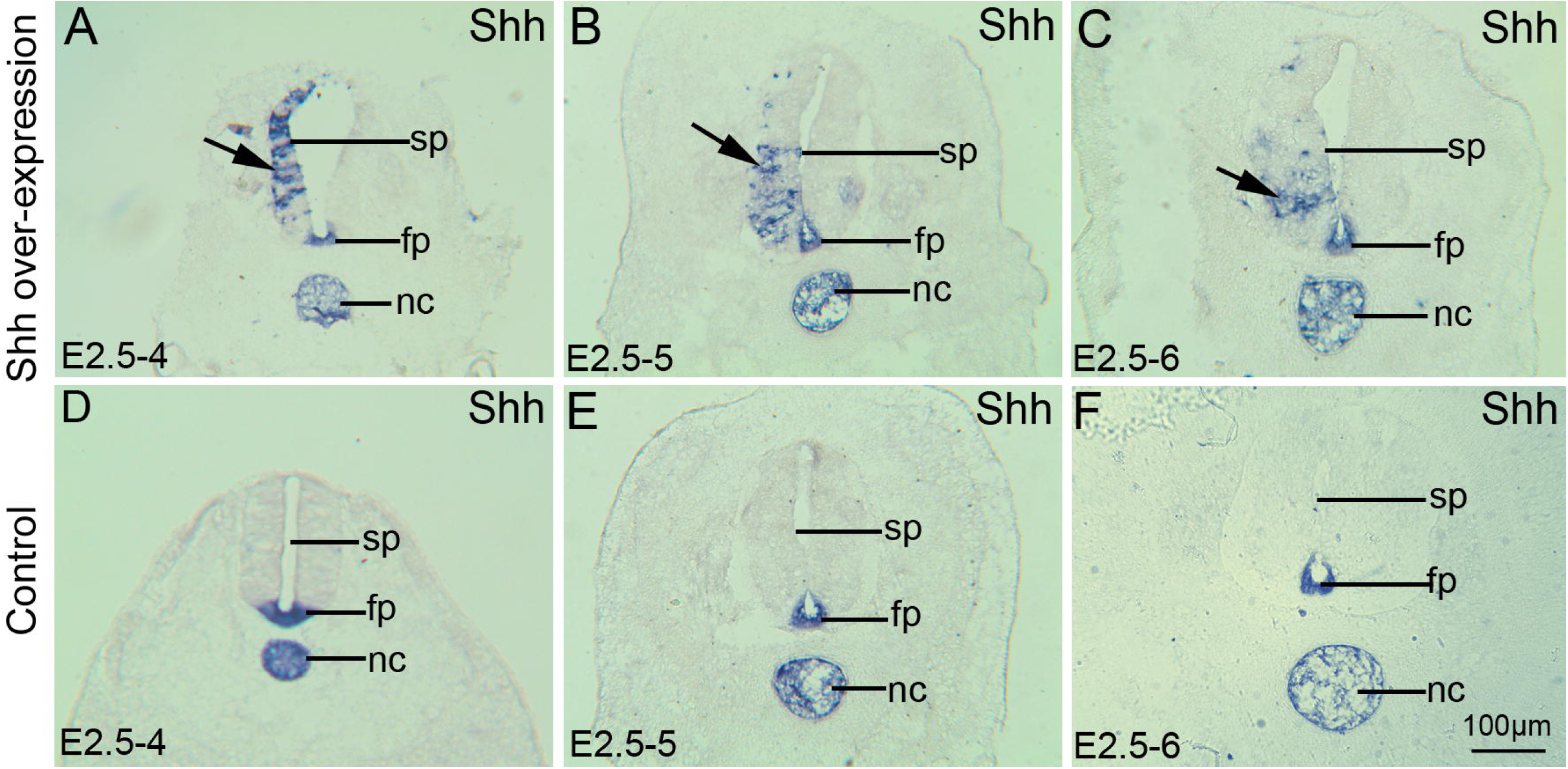
In situ hybridization demonstrates the overexpression of Shh. A-C: Shh overexpression following pCAGGS-Shh and pCAGGS-GFP co-transfection; D-F: Control group after pCAGGS-GFP transfection; A: at stage 24 (E4), B: at stage 27 (E5), C: at stage 28 (E6), D: at stage 24 (E4), E: at stage 27 (E5), and F: at stage 28 (E6), Arrows (→) indicate the areas of Shh overexpression. fp, floor plate; nc, notochord; sp, spinal cord; Scale bar = 100 μm in F for A-F.

### The effect of Shh over expression on microtubule-associated protein-2 (Map2) expression in motor column in the developing chicken spinal cord

Interestingly, MAP-2 labeling of motor columns following Shh overexpression in the spinal cord revealed structural abnormalities (Fig. 2A-L, arrow shown). In the control group, the structure of MAP-2 labeled motor column was normal (Fig. 2M-X). MAP-2 belongs to the microtubule-associated protein family. The proteins of this family are thought to participate in microtubule assembly, which is an essential step in neuritogenesis. MAP-2 isoforms are found predominately in neurons (Tucker, 1990). The principal functions of MAP-2 are to reduce the critical concentration of tubulin required to polymerize microtubules and to maintain neuronal morphology by regulating microtubule spacing (Caceres et al., 1992; Kalcheva et al., 1995). Even though MAP-2 is not a specific maker for motor neurons, motor neurons express MAP-2. Whether Shh overexpression affects the formation of the motor column by inhibiting the expression of MAP-2 in motor neurons is unclear. DAPI staining on the section slices showed a loss of cell nuclei on the transfected side of the motor column compared to non-transfected side (Fig. 2A-D, E-H). In the control group, the number of nuclei on the transfected side of the motor column was similar to that on the non-transfected side (Fig. 2M, Q). No GFP or MAP-2 positive neurons were observed in the motor column in the experimental group (Fig. 2 K-L). In the control group, GFP and MAP-2 positive neurons were observed in the motor column (Fig. 2 U-X). Therefore, it could be speculated that Shh may not inhibit the expression of MAP-2, but instead, modify the migration of motor neurons to the motor column. To verify this hypothesis, we used MNR2 to label motor neurons.

**Fig. 2.**
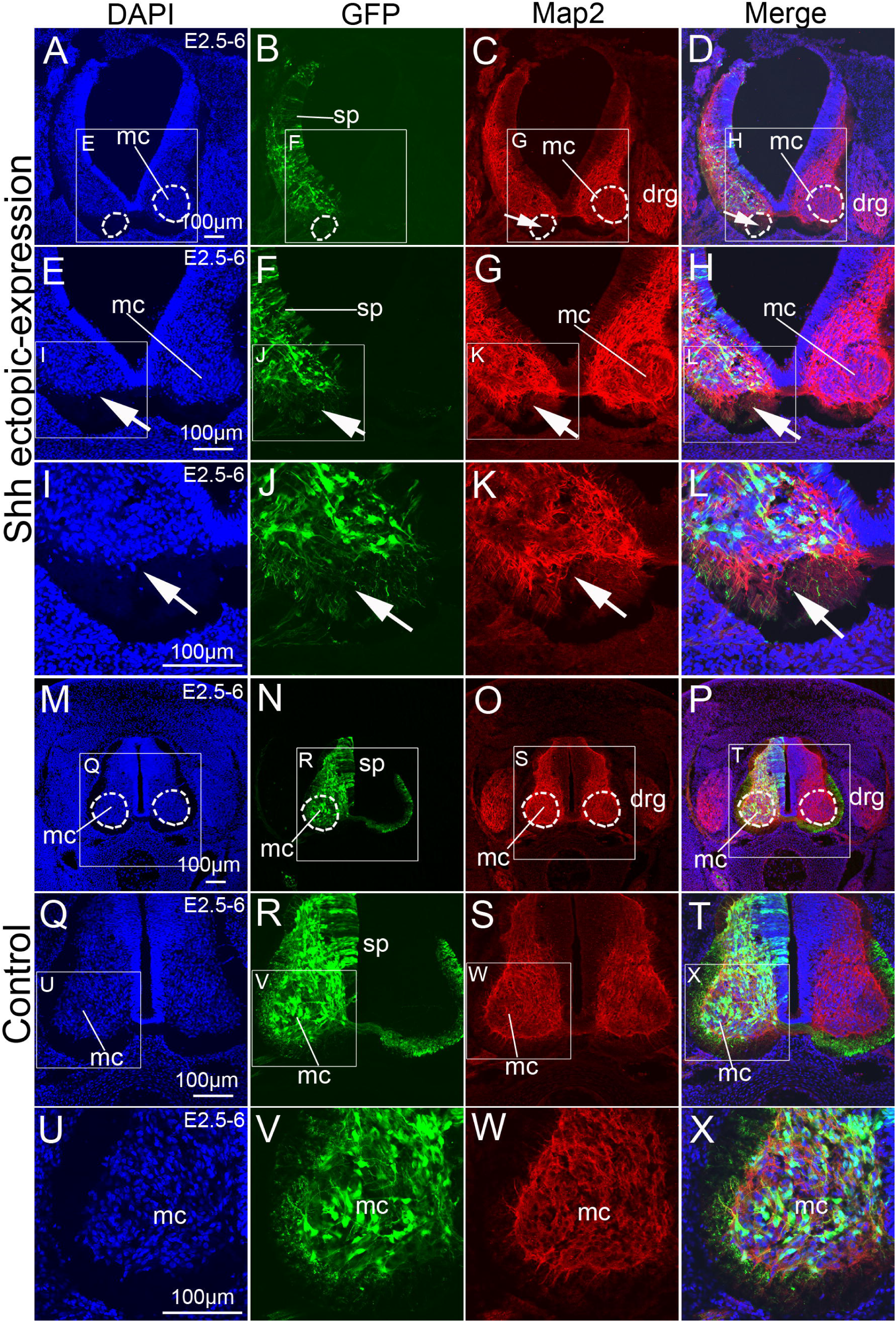
The effect of Shh overexpression on microtubule-associated protein-2 (Map2) expression within the motor column in the developing chicken spinal cord. A-L: Shh overexpression following pCAGGS-Shh and pCAGGS-GFP co-transfection at 84 h. DAPI nuclear staining (A, higher magnification of the ventral areas in E and I), GFP expression (B, higher magnification of the ventral areas in F and J, green), Map2 expression (C, higher magnification of the ventral areas in G and K, red), and merged images (D, higher magnification of the ventral areas in H and L). M-X: Control group after pCAGGS-GFP transfection at 84 h. DAPI nuclear stain (M, higher magnification of the ventral areas in Q and U), GFP expression (N, higher magnification of the ventral areas in R and V), Map2 expression (O, higher magnification of the ventral areas in S and W, red), and the merged image (P, higher magnification of the ventral areas in T and X). mc, motor column, sp, spinal cord. Arrows (→) indicate the areas of Map2 expression. Scale bars, 100 μm in A, E, I, M, Q, U for A-X, respectively.

### The effect of Shh overexpression on motor neuron (MNR2) positioning within the motor column in the chicken spinal cord

MNR2 is expressed selectively by motor neurons (MNs) in the developing vertebrate central nervous system. In order to investigate whether Shh affects the migration of motor neurons, or simply inhibits the expression of MAP-2 in motor neurons, we used MNR2 to specifically identify motor neurons. In the Shh overexpression group, MNR2 positive cells showed decreased accumulation in the motor column on the transfected side of the spinal cord as compared to the non-transfected side (Fig. 3A-H). However, in the control group, the distribution of MNR2 positive cells in the motor column on the transfected side was similar to that on the non-transfected side (Fig. 3I-P). Moreover, we observed morphological changes in the spinal cord with Shh overexpression (Fig. 3A-D). The spinal cord on the transfected side was curved outward, which was interpreted as the result of Shh overexpression rather than a physiological phenomenon (Fig. 3E-H). On the contrary, the morphology of the GFP-transfected side in the spinal cord was normal (Fig. 3I-L). In these areas, no outward curving was observed (Fig. 3M-P). Outward bending of the spinal cord in the areas of Shh overexpression has several potential explanations. It may be explained by the fact that Shh promotes proliferation of neuroepithelial cells, which leads to bending of the spinal cord outwards. In addition, these morphological changes may also be due to the effect of Shh on neuronal migration, especially that of motor neurons. The distribution of MNR2-labeled cells supports the effect of Shh on motor neuron migration. In order to investigate whether Shh can promote the proliferation of neuroepithelial cells, BrdU was used to label the proliferating cells.

**Fig. 3.**
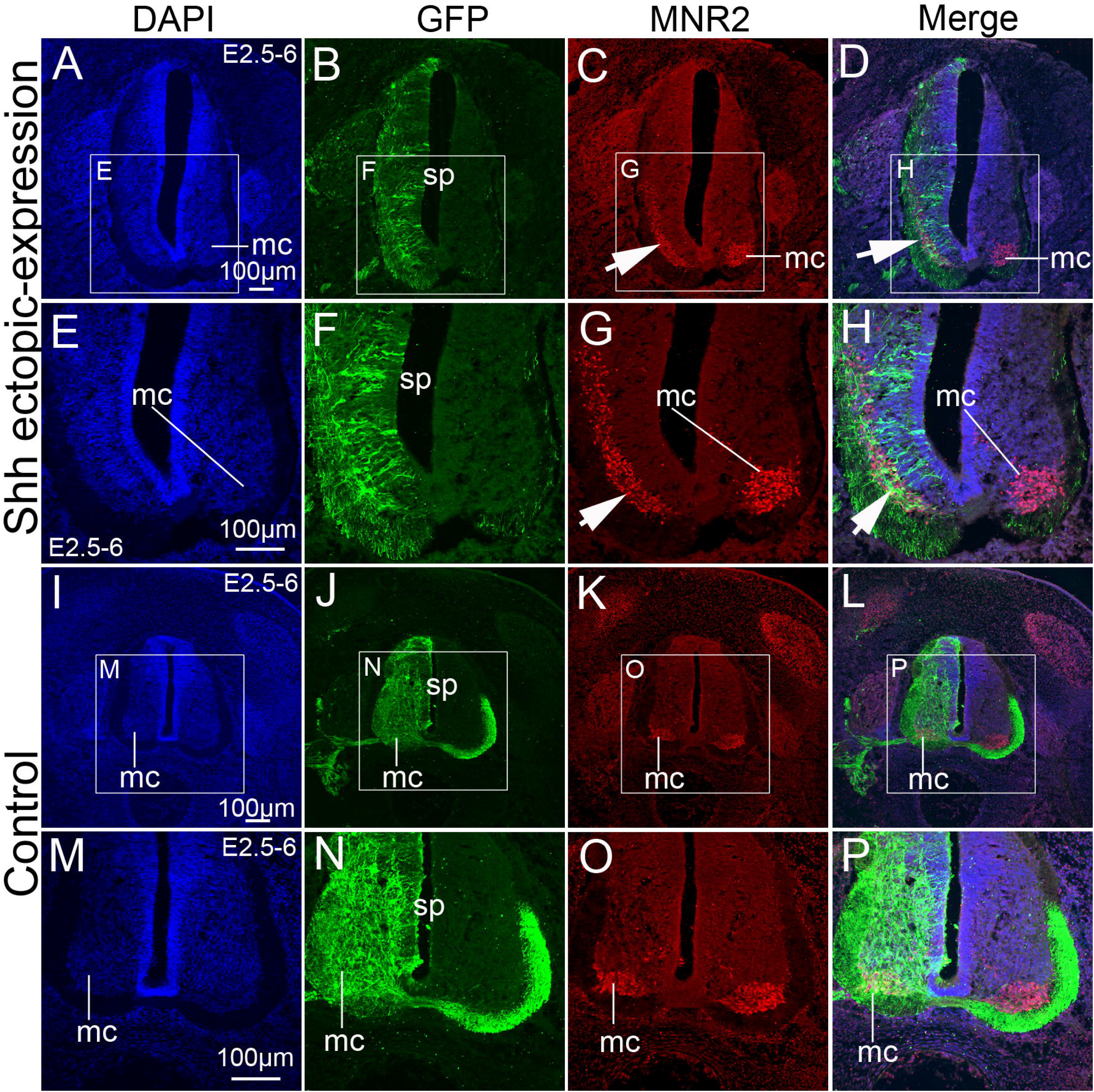
The effect of Shh overexpression on motor neuron (MNR2) positioning within the motor column in the chicken spinal cord. A-H: Shh overexpression following pCAGGS-Shh and pCAGGS-GFP co-transfection at 84 h. DAPI nuclear staining (A, higher magnification of the ventral areas in E), GFP expression (B, higher magnification of the ventral areas in F, green), MNR2 expression (C, higher magnification of the ventral areas in G, red), and merged images (D, higher magnification of the ventral areas in H). I-P: Control group after pCAGGS-GFP transfection at 84 h. DAPI nuclear stain (I, higher magnification of the ventral areas in M), GFP expression (J, higher magnification of the ventral areas in N), MNR2 expression (K, higher magnification of the ventral areas in O, red), and the merged image (L, higher magnification of the ventral areas in P). mc, motor column, Arrows (→) indicate the areas of MNR2 expression. Scale bars, 100 μm in A, E, I, M for A-P, respectively.

### The effect of Shh expression on neuroepithelial cell proliferation in the spinal cord during chicken embryo development

BrdU is a synthetic analog of thymidine commonly used for the detection of proliferating cells in living tissues (Lehner et al., 2011). BrdU was added 24 h before the spinal cord tissue was collected. Immunohistochemistry with anti-BrdU monoclonal antibody was used to reveal BrdU-positive cells. These cells were counted in the neural epithelium. The ratios of BrdU-positive cell number on the experimental (transfected) side over control (non-transfected) side were analyzed (Fig. 4a). Such a comparison between the experimental group versus control group (as shown in the Fig. 4a) indicated a significant increase in BrdU-positive cell numbers in the developing chick spinal cord, from stage 17 to 24, in Shh transfected tissue (Fig. 4A-D). The ratio of transfected to non-transfected side was 1.57±0.22 (n=3). In the control group, no difference in the number of BrdU-positive cells was observed between the GFP-transfected side and the non-transfected side of the spinal cord, from stage 17 to 24 (Fig. 4E-H). The ratio of transfected to non-transfected side was 1.12±0.14 (n=3). The ratios of transfected to non-transfected side were significantly different between the Shh overexpression group and control group (p<0.01, Fig. 4b). A comparison between the side of the spinal cord transfected with Shh and the control non-transfected side (as shown in the Fig. 4c) indicated a significant decrease in BrdU-positive cells in the developing chick spinal cord, from stage 17 to 29 (Fig. 4I-L). The ratio of transfected side to non-transfected side was 0.70±0.32 (n=3). In the control group, no difference in the number of BrdU-positive cells between the transfected and non-transfected side in the spinal cord was observed from stage 17 to 29 (Fig. 4M-P). The ratio of transfected side to non-transfected side was 0.98±0.19 (n=3). The ratios of transfected side to non-transfected side in Shh overexpression group versus control group were significantly different (p<0.01, Fig. 4d). The decrease in the number of BrdU-labeled cells on the side with Shh overexpression compared to the contralateral side was particularly visible in the ventral areas of the spinal cord (Fig. 4Q-T). The ratio of transfected side to non-transfected side was 0.53±0.27 (n=3). In the control group, no differences were observed (Fig. 4U-X). The ratio of transfected side to non-transfected side was 1.17±0.11 (n=3). As shown in Fig. 4e, the ratio of the number of BrdU-positive cells on the transfected side to that on the non-transfected side was significantly different in the Shh overexpression group compared to control group (p<0.01, Fig. 4f). Interestingly, in stage 24 (E4), Shh promoted neuroepithelial cell proliferation (Fig. 4b), while in stage 26 (E6) it had an inhibitory effect (Fig. 4d, f). Therefore, we speculated that Shh not only affects the proliferation of neural precursor cells but also their differentiation. It is possible that, in response to Shh misexpression in early stages of embryonic development, neural precursor cells were induced to differentiate into nerve cells therefore losing their ability to proliferate, which is why the number of proliferating cells was significantly decreased compared to control group.

**Fig. 4.**
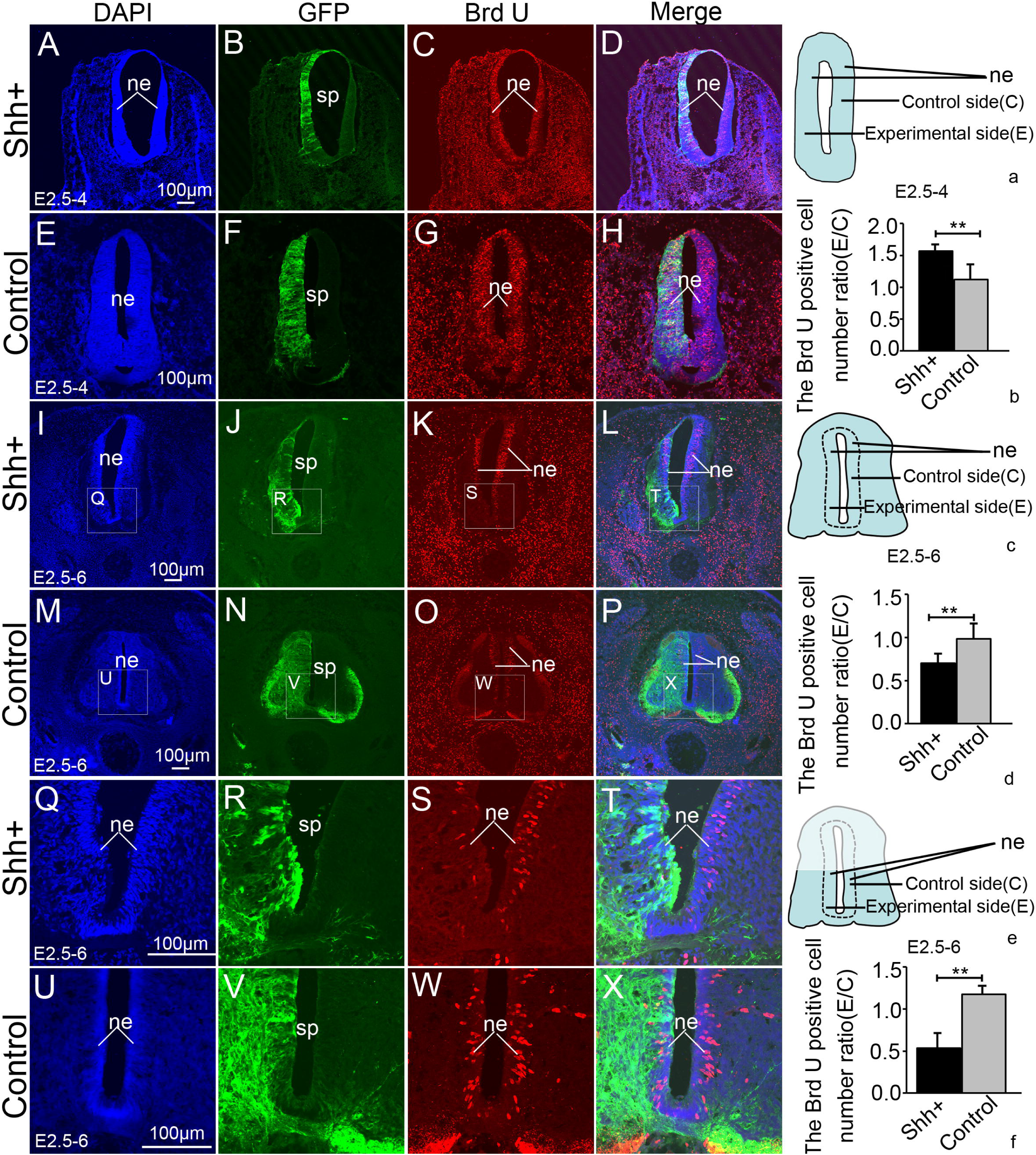
The effect of Shh overexpression on neuroepithelial cell proliferation in the spinal cord during chicken embryo development. A-D: Shh overexpression following pCAGGS-Shh and pCAGGS-GFP co-transfection for 36 h. DAPI nuclear staining (A), GFP expression (B, green), BrdU expression (C, red), and merged images (D). E-H: Control group after pCAGGS-GFP transfection at 36 h. DAPI nuclear stain (E), GFP expression (F), BrdU expression (red; G), and merged image (H). I-L: Shh overexpression following pCAGGS-Shh and pCAGGS-GFP co-transfection at 84 h. DAPI nuclear staining (I, higher magnification of the ventral areas in Q), GFP expression (J, higher magnification of the ventral areas in R, green,), BrdU expression (K, higher magnification of the ventral areas in S, red), and merged images (L, higher magnification of the ventral areas in T). M-P: Control group after pCAGGS-GFP transfection at 84 h. DAPI nuclear stain (M, higher magnification of the ventral areas in U), GFP expression (N, higher magnification of the ventral areas in V), BrdU expression (O, higher magnification of the ventral areas in W, red), and merged image (P, higher magnification of the ventral areas in X). a, the pattern of spinal tissue slice section at 36 h. b, the ratio of BrdU-positive cell numbers on the experimental side to those on the control side (E/C) at 24 h; c, the pattern of spinal tissue slice section at 84 h. d, the ratio of BrdU-positive cell numbers on the experimental side to those on the control side (E/C) at 84 h; e, the pattern of spinal tissue slice section in ventral areas at 84 h. f, the ratio of BrdU-positive cell numbers on the experimental side to those on the control side (E/C) in ventral areas at 84 h. Data are presented as mean ± S.D. **p<0.01. ne, neuroepithelial cells. Scale bars, 100 μm in A, E, I, M, Q, U for A-X, respectively.

### The effect of Shh overexpression on Pax3 and Pax7 expression in the spinal cord during chicken embryo development

Shh affects not only the differentiation and proliferation of ventral cells, but also the expression of dorsal genes during chicken embryonic development. The expression of the nuclear proteins Pax3 and Pax7 was therefore investigated. The results showed that Pax3 expression was inhibited at the side of Shh overexpression position compared to the control, non-transfected, side (Fig. 5A-F, arrow), which suggests that early expression of Shh inhibits Pax3 expression. However, no differences in the expression between the two sides of the spinal cord were observed in the control group (Fig. 5G-L). Further, the mean optical density ratios of the experimental (transfection) side to the control (no transfection) side were analyzed (Fig. 5a). Cell numbers in the control group were significantly (p<0.01) higher than those in the Shh overexpression group (Fig. 5 b). Pax7 expression was also inhibited at the side with Shh overexpression position compared to the control non-transfected side (Fig. 5M-R, arrow). No differences in Pax7 expression were observed between the transfected versus non-transfected side in the control group (Fig. 5S-X). The numbers of Pax7-positive cells in the control group were significantly (p<0.01) higher than in the Shh overexpression group (Fig. 5c). Additionally, the percentage of commissural axons projecting to the contralateral side in the Shh overexpression group was significantly lower in comparison to the control (Fig. 5d, p<0.01). Therefore, our results suggest that Shh overexpression may inhibit the commissural axons projecting to the contralateral side in the spinal cord during chicken embryo development.

**Fig. 5.**
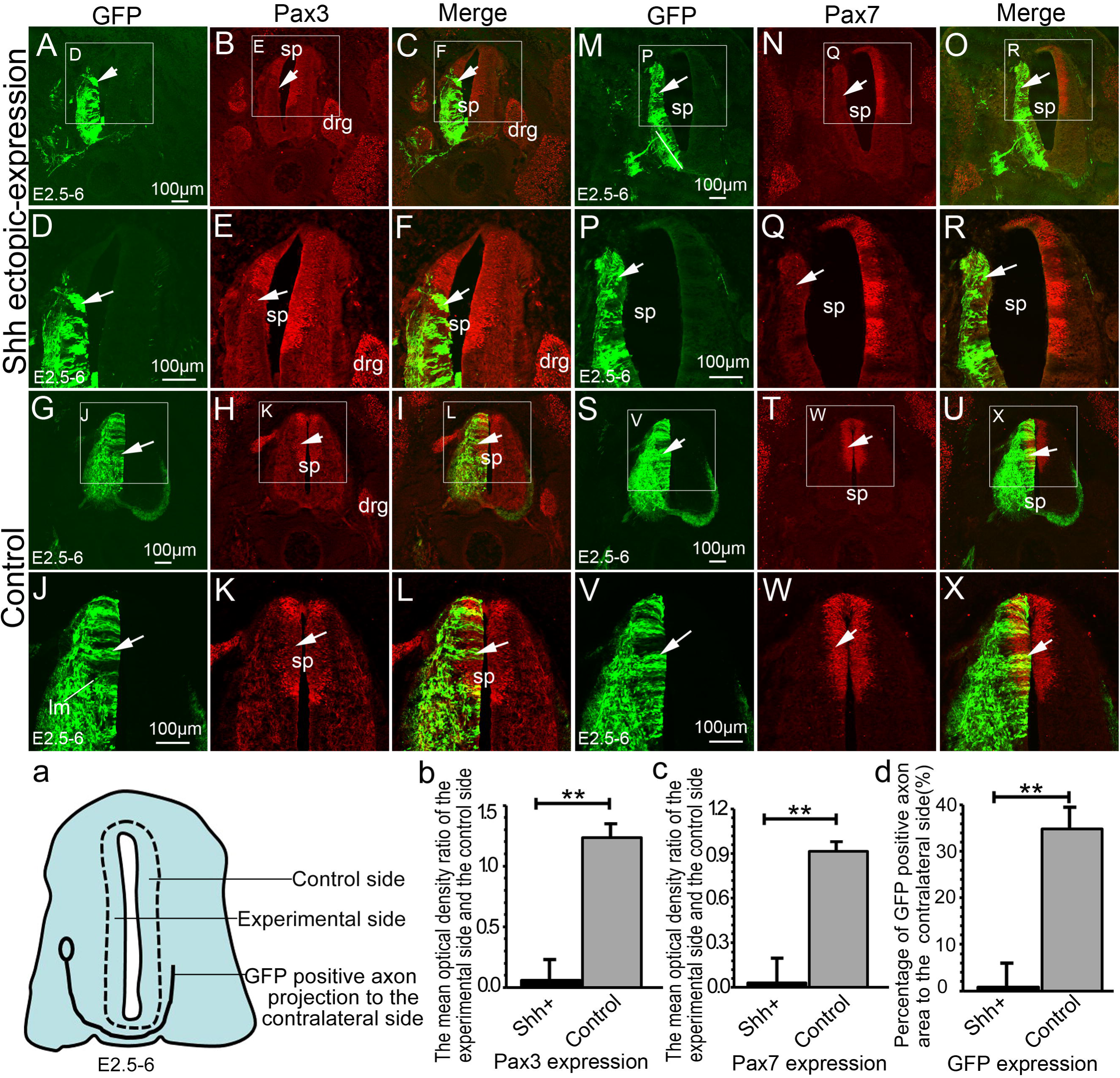
The effect of Shh overexpression on Pax3 and Pax7 expression in the spinal cord during chicken embryo development. A-F: Shh overexpression group with pCAGGS-Shh and pCAGGS-GFP plasmid co-transfection, showing GFP (A, higher magnification in D), Pax3 (B, higher magnification in E) expression, and merged image (C, higher magnification in F). G-L: Control group with pCAGGS-GFP plasmid transfection, showing GFP (G, higher magnification in J), Pax3 (H, higher magnification in K) expression, and merged image (I, higher magnification in L). M-R: Shh overexpression group with pCAGGS-Shh and pCAGGS-GFP plasmid co-transfection, showing GFP (M, higher magnification in P), Pax7 (N, higher magnification in Q) expression, and merged image (O, higher magnification in R). S-X: Control group with pCAGGS-GFP plasmid transfection, showing GFP (S, higher magnification in V), Pax7 (T, higher magnification in W) expression, and merged image (U, higher magnification in X). a, the pattern of spinal tissue slice section. b, the mean optical density ratio of the experimental side to the control side; c, the mean optical density ratio of the experimental side to the control side; d, percentage of GFP positive axonal area projecting to the contralateral side (%). Data are presented as mean ± S.D. **p<0.01. drg, dorsal root ganglion; sp, spinal cord. Arrows (→) indicate the areas of Pax3 or Pax7 expression. Scale bars, 100 μm in A, D, G, J for A-L. 100 μm in M, P, S, V for M-X, respectively.

To assess the morphological changes that are induced by shh overexpression along the transfected spinal cords, the rostro-caudal series of sections were obtained. These series sections result confirmed that the shh overexpression perturbed axon projections (Fig 6). The shh overexpression lead to commissural axons projecting to the contralateral side weak mlc and almost no ilc (Fig. 6A-G), as H shows. In the control there have normal commissural axons projecting to the contralateral side and axons arrived to mlc and ilc (Fig. 6I-O), as Fig 6P shows.

**Fig. 6.**
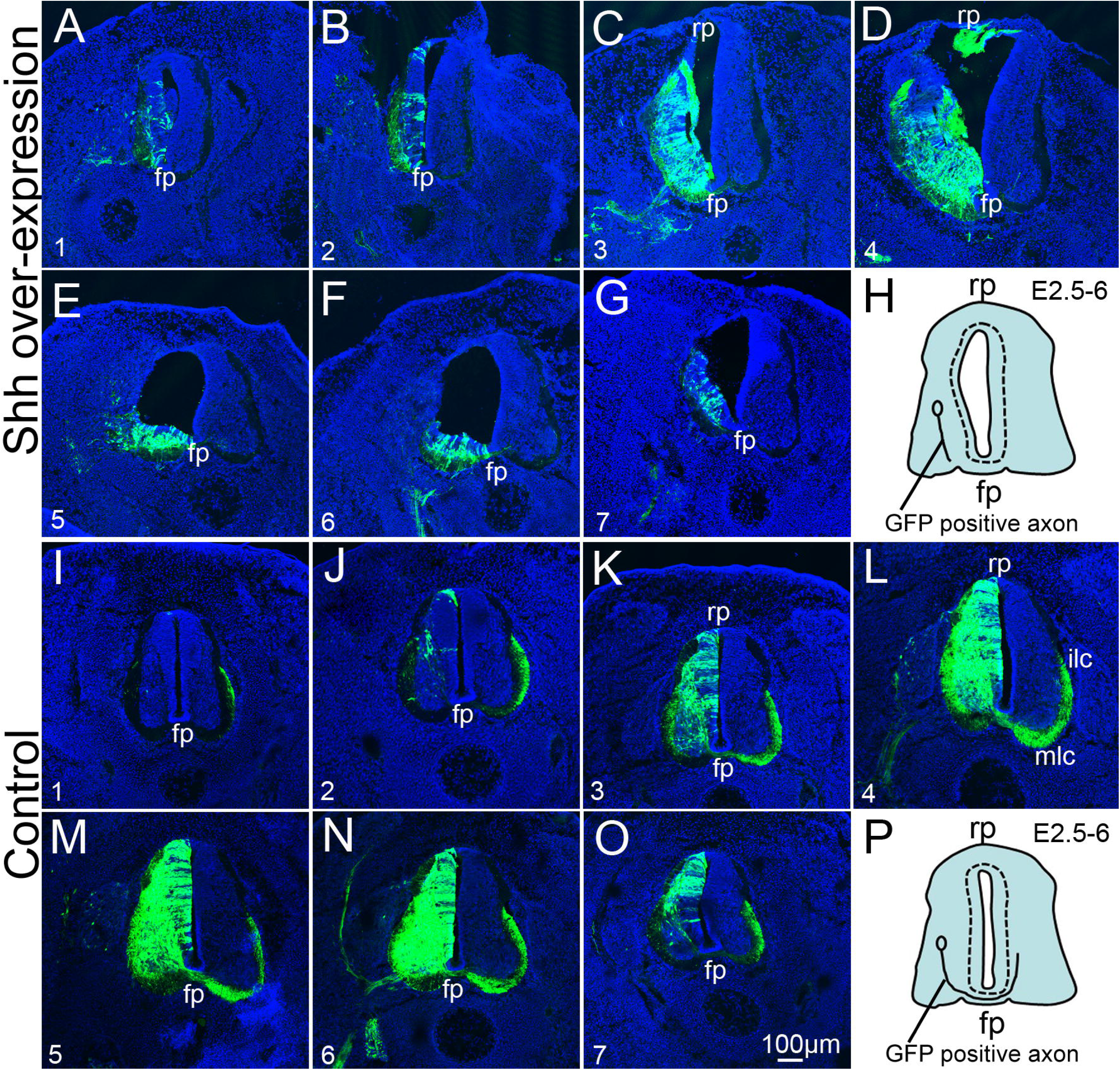
The effect of shh overexpression on commissural axons projection in the spinal cord during chicken embryo development. Rostro-caudal series of transverse sections after electroporation of the shh overexpression (shh overexpression; A-G); H, the pattern of commissural axons projection. The GFP alone expression (control; I-O); P, the pattern of commissural axons projection. In ovo electroporation was performed at E2.5 (2.5 days’ incubation) and the positive embryos were collected at E6. Abbreviations: fp, floor plate; ilc, intermediate longitudinal commissural axons; mlc, medial longitudinal commissural axons; rp, roof plate. Scale bar: 100μm in O for A-G and I-O.

## DISCUSSION

Shh is one of three proteins in the mammalian hedgehog signaling family, the others being desert hedgehog and Indian hedgehog. Shh is the most studied hedgehog signaling pathway. It plays a critical role in the patterning of vertebrate embryonic nervous system, including the brain and the spinal cord, during development (Chiang et al., 1996). Shh is a secreted protein that mediates signaling activities in the notochord and the floor plate (Patten and Placzek, 2000). One of early functions of the notochord is to induce differentiation of ventral cell types, such as floor plate cells and motor neurons in the overlying neural ectoderm (Chiang et al., 1996). Shh is considered to play an important role during spinal cord development (Martí et al., 1995), given its predominant expression in the notochord and floor plate during embryo development. In this study, we demonstrate that the overexpression of Shh affects motor neuron positioning in the spinal cord during chicken embryo development.

Our results show that Shh overexpression affects the expression pattern of MAP-2. The abnormal pattern of MAP-2 expression was not due to the inhibition by Shh, but to the effect of Shh on the migration of motor neurons, preventing them from reaching their positions within the motor column accurately. In order to confirm these results, MNR2 labeling was performed to precisely identify motor neurons. The results showed that, in the Shh overexpression group, the transfected side showed a deficit in the accumulation of MNR2 positive cells in the motor column compared to non-transfected side. These results were consistent with the expected ones, indicating that the Shh overexpression, rather than inhibiting the expression of MAP-2, affected the migration of motor neurons, which led to the absence of MAP-2 expression in the motor column region. Besides Shh, factors such as HB9, Islet1, Islet2, Lim1, and Foxp1, if misexpressed, could also induce defective motor neuron positioning within the spinal cord during embryo development (Bréjot et al., 2006; Odden et al., 2002; Hutchinson and Eisen, 2006; Kania et al., 2000; Otaegi et al., 2011; Rousso et al., 2008). The mechanisms by which different molecules affect the migration of motor neurons are different. Shh acts in a graded fashion to pattern the dorsal-ventral axis of the vertebrate spinal cord. This is a dynamic process in which increasing concentrations and the duration of exposure to Shh generate neurons with successively more ventral identities (Ribes and Briscoe, 2009). Thus, Shh ligand secreted by the notochord induces distinct ventral cell identities in the adjacent spinal cord by a concentration-dependent mechanism (Chamberlain et al., 2009). Normally, the concentration of Shh increases gradually from dorsal to ventral regions. The highest concentrations of the Shh ligand are found in the most ventral regions of the neural tube and notochord, while lower concentrations are found in the more dorsal regions of the neural tube (Ribes et al., 2009). In our experiment, the overexpression of Shh in the spinal cord induced structural abnormalities in the nerve column. One possibility is that motor neurons failed to migrate to the right position, possibly due to high concentrations of Shh while another is that the structural abnormalities are a result of the inhibition of neuroepithelial cell differentiation into motor neurons.

Therefore, in this study, the proliferation of neuroepithelial cells was investigated using labeling with BrdU. BrdU is incorporated into newly synthesized DNA in the replicating cells during the S phase of the cell cycle, as a substitute for thymidine. Antibodies specific for BrdU can be used to detect this compound incorporated into the DNA, thus indicating the cells with actively replicating DNA during BrdU administration (proliferating cells). The results of our study show that cell proliferation in the early stage (stage 18-24, E3-E4) was higher than in the late stage (stage 27-29, E5-E6). Does this mean that the overexpression of Shh promotes the proliferation of neuroepithelial cells in the early stage (stage 18-24, E3-E4), but inhibits proliferation in the late stage (stage 27-29, E5-E6)? Studies have shown that Shh acts in a concentration-dependent manner (Placzek et al., 1990) so that lower concentrations of Shh promote cellular proliferation and induction of various ventral neural cell types (Ericson et al., 1996), while high concentrations of Shh inhibit cellular proliferation (Wilson and Maden, 2005). We believe that promoting cell proliferation is only one of the effects of Shh, the other one being the promotion of neural precursor cell differentiation. In the early stage (stage 18-24), Shh promotes the neural precursor cell differentiation into neurons, and these neurons then lose the ability to proliferate. Shh affected the formation of motor neurons by inducing a defect in their migration to the motor column and, consequently, altering their distribution in the border of the gray matter and leading to the formation a band. The ultimate result was the abnormal structure of the motor column. The effect of Shh on neural precursor cell differentiation requires further research.

It is thought that Shh gradient determines multiple different cell fates by a concentration and time-dependent mechanism that induces the expression of several transcription factors in ventral progenitor cells (Chamberlain et al., 2008). In this study, we examined the expression of dorsal transcription factors Pax3 and Pax7. Our results showed that the expression of Pax3 and Pax7 was inhibited in the regions of Shh overexpression. Pax3 and Pax7 participate in the sonic hedgehog (Shh) signaling pathway and are inhibited by Shh overexpression (Lin et al., 2016). Our previous study indicated that the transcription factors Pax3 and Pax7 play important roles in regulating morphogenesis and cell differentiation in the developing spinal cord (Lin et al., 2016). Shh has also been shown to act as an axonal guidance molecule. Studies have demonstrated that Shh attracts commissural axons at the ventral midline of the developing spinal cord (Charron et al., 2003). In this study, we also showed that the overexpression of Shh significantly inhibited the commissural axons from projecting to the contralateral side. Our previous study indicated that the transcription factors Pax3 play important roles in induces cell aggregation and perturbs commissural axon projection during embryonic spinal cord development (Lin et al., 2017). In this study showed that the shh overexpression inhibited the expression of Pax3 and Pax7. Therefore, the effect of Shh on the commissural axon projection may be related to the effects of Shh on the expression of Pax3 and Pax7.

The current study provided evidence that Shh affects motor neuron positioning in the spinal cord during chicken embryo development. The overexpression of Shh in the spinal cord not only altered the positioning of the motor neurons, but also resulted in the abnormal structure of the motor column. At the same time, Shh misexpression inhibited the expression of genes related to the dorsal development and further perturbed commissural axon projections during chicken embryo development.

## MATERIALS AND METHODS

### Embryo and tissue preparation

Fertilized eggs of Sea blue brownhad, obtained from a local farm (HWS-150, JingHong, China) were incubated at 37.8 °C and 65% humidity. The Hamburger and Hamilton (Hamburger V and Hamilton HL, 1992) system was used to stage the embryos. The embryos were studied at stage 18 (E2.5) to stage 29 (E6), with at least three embryos at each stage.

### In ovo electroporation

The Shh overexpression plasmid was a gift by Redies (Prof. Christoph Redies, Institute of Anatomy I, Jena University Hospital, Teichgraben 7, D-07743 Jena, Germany). The plasmid pCAGGS-GFP (green fluorescent protein) was derived by our laboratory. All plasmids used were extracted with a kit (Cwbio, Beijing. China) and diluted in water.

The in ovo electroporation protocol was modified from our previous publications (Luo et al., 2006; Yang et al., 2015; Lin et al., 2016). A stereomicroscope was used in all the steps of the procedure. In brief, fertilized eggs were incubated until stage 18 (E2.5). Then, 3-4 mL of albumin was removed from the egg without disrupting the yolk. Further, an incision into the shell was performed carefully using a pair of curved scissors to obtain a 1-2-cm diameter window without touching the embryo. A mixture of 4 μg/μL of pCAGGS-Shh, 0.25 μg/μL of pCAGGS-GFP plasmid, and Fast Green dye (0.01%) or that of 0.25 μg/μL of pCAGGS-GFP plasmid as a control and Fast Green dye (0.01%), were injected and loaded into the neural tube lumen using a mouth pipette until the dye filled the entire space. The electrodes were then immediately placed on both sides of the embryonic neural tube in parallel. A total of six 18-volt pulses, which lasted for 60 ms, with a pause of 100 ms between each pulse, were delivered. Bubbles near the electrodes indicated that the technique was successfully performed. After the electroporation (CUY-21, Nepa Gene, Japan), the electrodes were carefully removed, and the egg was sealed with a tape. The treated eggs were then placed back in the incubator until they reached the desired stage for sample collection, fixation, and analysis. For bromodeoxyuridine (BrdU) labeling, 5 μg/μL of BrdU was added into the embryo 24 hours prior to fixation.

### Tissue section

When the embryos were at E6 (stage 26), samples of the spinal cord tissue were collected. The embryos were fixed in 4% formaldehyde solution for 6-24 h, depending on their size. After fixation, the tissue was immersed in 18% sucrose solution, embedded in Tissue-Tec O.C.T. compound (Sakura Finetek, USA), frozen in liquid nitrogen, and stored at -80 °C until required. Samples were sectioned using a cryotome (Leica 1850, Germany) and 20-μm thick sections were mounted on Poly-L-lysine coated slides.

### cRNA probe synthesis and in situ hybridization

Digoxigenin-labeled sense and antisense cRNA probes were transcribed in vitro using purified PBS-SK plasmids containing the full-length Shh according to the manufacturer’s instructions (Roche, Germany). Sense cRNA probes were used as negative controls for in situ hybridization.

For in situ hybridization, 20-μm thick cryosections were fixed with 4% formaldehyde in PBS and pretreated with proteinase K and acetic anhydride. The sections were hybridized overnight with a cRNA probe at a concentration of about 3 ng/μL at 70^o^C in hybridization solution (50% formamide, 3⨯ SSC, 10 mM EDTA, 10% dextran sulfate, 1⨯ Denhardt’s solution, 42 μg/mL yeast transfer RNA and 42 μg/mL salmon sperm DNA; Roche, Germany). The sections were washed to remove unbound cRNA by RNase reaction, and then incubated with alkaline phosphatase-coupled anti-digoxigenin Fab fragments (Roche, Germany) at 4^°^C overnight. For the visualization of labeled mRNA, a substrate solution of nitroblue tetrazolium salt (NBT, Fermentas, Lithuania) and 5-bromo-4-chloro-3-indoyl phosphate (BCIP, Fermentas, Lithuania) were added.

### Immunohistochemistry

For immunohistochemistry, sections were fixed with 4% paraformaldehyde in PBS for 15 min at 37^o^C. Following another TBS wash, a blocking solution (2% sheep serum, 4% bovine serum albumin, 0.3% Triton X-100, and 0.1% sodium azide in Tris-buffered saline, TBS, sheep serum and bovine serum albumin; Beijing Dingguo co. LTD, China) was applied to tissue sections, for 1 h at room temperature. The primary antibodies were then applied overnight at 4^o^C. The primary antibodies used in the present study were rabbit anti Map2 polyclonal antibody (Abcam, United Kingdom, 1:500 dilution), mouse anti chicken MNR2 monoclonal antibody (DSHB, USA; 1:100 dilution), mouse anti Pax3 monoclonal antibody (DSHB, USA; 1:100 dilution), mouse anti Pax7 monoclonal antibody (DSHB, USA; 1:100 dilution), and mouse anti BrdU monoclonal antibody (ZSGB-BIO, China; 1:100 dilution). For BrdU detection, sections were incubated in 2 N HCl for 30 minutes followed by 0.1 M Na_2_B_4_O_7_ (pH 8.5), and then rinsed several times in TBS before the incubation with anti-BrdU. Next, the appropriate goat-anti-rabbit Cy3-labeled (Jackson Immuno Research, Europe Ltd, 1:1000 dilution), goat-anti-mouse Cy3-labeled (Jackson Immuno Research, Europe Ltd, 1:1000 dilution) or goat-anti-rabbit FITC-labeled (ZSGB-BIO, China; 1:100 dilution) secondary antibodies were applied for 2 h at 25^o^C. A similar process was employed for double staining. Finally, DAPI (4’,6-diamidino-2-phenylindole, DAPI, Roche, Germany) was used to stain all cell nuclei.

### Antibody characterization

See Table 1 for a list of all antibodies used. This study were used antibodies has been described extensively. The staining patterns produced by all of the antibodies were similar to those described previously.

**Table 1.** Antibodies Used in This Study

| Antigen | Immunogen | Detail of antibodies | Working dilution |
| --- | --- | --- | --- |
| **Pax3** | Chicken PAX3 (aa 298-481), recombinant protein made in E. coli | DSHB, USA, (PAX3), mouse, monoclonal antibody, IgG2a | 1:100 |
| **Pax7** | Chicken PAX7 (aa 352-523), recombinant protein made in E. coli | DSHB, USA, (PAX7), mouse, monoclonal antibody, IgG1 | 1:100 |
| **MNR2** | C-terminal portion of chicken MNR2-GST fusion protein made in E. coli | DSHB, USA, (81.5C10) mouse, monoclonal antibody, IgG1 | 1:100 |
| **Map2** | Rat microtubule associated protein 2 | Abcam, United Kingdom, (ab32454), Rabbit, polyclonal | 1:500 |
| **BrdU** | BrdU ( Bromodeoxyuridine ) | ZSGB-BIO, China, (ZM-0013), mouse, monoclonal antibody, IgG1 | 1:100 |

Mouse monoclonal antibody against Pax3 was obtained from the Developmental Studies Hybridoma Bank (catalogue no. pax3, RRID:AB_528426, mouse, monoclonal antibody, IgG2a). To generate this antibody, the cDNA region that corresponded to amino acids 298-481 of the C.terminal region of quail Pax3 was cloned by PCR into the E. coli expression vector (Joven et al., 2013). Venters et al. (2004) confirmed that the Pax3 antibody stains an single band of about 60 kDa on Western blots of extract from E3 chicken neural tube and notochord. In addition, the expression pattern of Pax3 protein obtained in the present study was similar to that reported previously for Pax3 protein (Lin et al., 2017).

The mouse monoclonal antibody against Pax7 (DHSB, Catalogue No. pax7, RRID: AB_528428) was generated by Dr. Atsushi Kawakami (Kawakami et al., 1997). The DNA region corresponding to amino acids 352-523 of chick Pax7 was cloned by PCR into the E. coli expression vector (Joven et al., 2013). Anti Pax7 antibody detects three bands on Western blots of chicken brain tissue (Ferran et al., 2009). The staining pattern of the Pax7 antibody obtained in the present study was the same as that reported previously (Kobayashi et al., 2013; Lin et al., 2016).

The specificity of anti-MNR2 (81.5C10) was determine by comparison of the labeling patterns obtained by immunohistochemistry and by in situ hybridization in the chick embryo spinal cord (Tanabe et al., 1998). The staining pattern of the antibody in the this study was consistent with previous reports (Tanabe et al.,1998; Kobayashi et al., 2013).

Anti-MAP2 was raised against rat microtubule-associated protein 2. Synthetic peptide conjugated to KLH derived from within residues 1-100 of Rat MAP2. Anti MAP2 antibody detects two bands on Western blots of mouse brain tissue lysate total protein (260,280 kDa). Additional bands at: 110 kDa,199 kDa,65 kDa are unsure as to the identity of these extra bands (Sigma Product Sheet).

Monoclonal mouse anti-BrdU IgG1 (1:100; ZSGB-BIO, China, (ZM-0013), mouse, monoclonal antibody, IgG1), recognizes BrdU. This antibody reacts with BrdU incorporated into single-stranded DNA, attached to a protein carrier and free BrdU.

### Microscopy

The whole embryo was imaged under a stereo fluorescence microscope (LEICA M205FA, Germany) equipped with a digital camera (LEICA DFC425C, Germany). In situ hybridization sections were viewed under a microscope (Nikon ECLIPSE 80i, Japan), which was equipped with a digital camera (LEICA DFC300FX, Germany). Immunohistochemistry sections were imaged under a confocal microscope (Olympus ix81, Japan).

### Statistical analysis

The average optical density and area of fluorescence intensity were calculated using Plus Image-Pro 6 software (Media Cybernetics, USA), and the data were analyzed by Statistics 17.0 SPSS software (IBM, USA). All data are presented as the mean ± standard deviation (S.D.), of at least three independent experiments. The significance of differences among the transfection groups was determined using ANOVA, where p-value <0.05 was considered as significant.

## Author contributions

Conceived and designed the experiments: Juntang Lin. Performed the experiments: Ciqing Yang, Xiaoying Li, Qiuling Li, Qiong Li, Bichao Zhang. Analyzed the data: Han Li, Ciqing Yang. Wrote the paper: Ciqing Yang.

